# Using BERT to identify drug-target interactions from whole PubMed

**DOI:** 10.1101/2021.09.10.459845

**Authors:** Jehad Aldahdooh, Markus Vähä-Koskela, Jing Tang, Ziaurrehman Tanoli

## Abstract

**Background:** Drug-target interactions (DTIs) are critical for drug repurposing and elucidation of drug mechanisms, and they are collected in large databases, such as ChEMBL, BindingDB, DrugBank and DrugTargetCommons. However, the number of studies providing this data (~0.1 million) likely constitutes only a fraction of all studies on PubMed that contain experimental DTI data. Finding such studies and extracting the experimental information is a challenging task, and there is a pressing need for machine learning for the extraction and curation of DTIs. To this end, we developed new text mining document classifiers based on the Bidirectional Encoder Representations from Transformers (BERT) algorithm. Because DTI data intimately depends on the type of assays used to generate it, we also aimed to incorporate functions to predict the assay format.

**Results:** Our novel method identified and extracted DTIs from 2.1 million studies not previously included in public DTI databases. Using 10-fold cross-validation, we obtained ~99% accuracy for identifying studies containing drug-target pairs. The accuracy for the prediction of assay format is ~90%, which leaves room for improvement in future studies.

**Conclusion:** The BERT model in this study is robust and the proposed pipeline can be used to identify new and previously overlooked studies containing DTIs and automatically extract the DTI data points. The tabular output facilitates validation of the extracted data and assay format information. Overall, our method provides a significant advancement in machine-assisted DTI extraction and curation. We expect it to be a useful addition to drug mechanism discovery and repurposing.

## BACKGROUND

The average cost of developing a new drug ranges in billions of dollars, and it takes about 9–12 years to bring a new drug to the market [1]. Hence, finding new uses for already approved drugs is of major interest to the pharmaceutical industry. This practice, termed drug repositioning or drug repurposing, is attractive because of its potential to speed up drug development, reduce costs, and provide treatments for unmet medical needs [2]. Central to drug discovery and repositioning are drug-target interactions (DTI), meaning the qualitative, quantitative, and relative interactions of compounds with the molecules that regulate cellular functions. DTIs are catalogued in public databases, which classify DTIs as binary (contains both active and inactive interactions), unary (only active interactions) or as quantitative (in terms of IC50, Kd, Ki etc.) [3]. The most well-known resources for quantitative bioactivity interactions are ChEMBL [4], BindingDB [5], PubChem [6], GtopDB [7] and DrugTargetCommons [8][9]. These resources contain experimental data for millions of compounds across thousands of protein targets. Still, the combined non-overlapping studies in these databases are less than 0.1 million and cover around 3,000 protein targets with an average of 7.33 interactions per target [10]. Moreover, each data resource focuses only on specific journals for data curation. For instance, ChEMBL and DrugTargetCommons primarily focus on Medicinal chemistry, Nature biotechnology and a few other journals. However, there are more than 7000 journals and 32M studies on PubMed [11]. Large fraction of the skipped studies in drug-target databases may contain experimentally tested DTIs. However, mining PubMed manually is not efficient. Therefore, there is a need to develop Natural Language Processing (NLP) based semi-automated text classifiers that can initially identify relevant studies (called ‘triage’) and dig further to extract actual DTIs reported in the full text of the articles.

Text classification is a well-known problem in NLP. The objective is to assign predefined categories to a given text sequence (in this case, it could be an abstract, title or full text for the study). One of the pre-processing step is to map textual data into numeric features [12] to make it understandable by the classification model. Mapping of textual information into numeric features can be performed using pre-trained models on a large corpus of texts. Pre-trained language models on large text corpora are proven to be adequate for the task of text classification with a decrease in computational costs at runtime [13]. Among those are the word embedding based models, such as word2vec [14] and GloVe [15] and contextualized word embedding models, such as CoVe [16] and ELMo [17]. Others are sentence-level models, such as ULMFiT [18]. More recently, pre-trained language models are shown to be helpful in learning common language representations by utilizing a large amount of un-labelled data: e.g., OpenAI GPT [19] and BERT [20]. Bidirectional Encoder Representations from Transformers (BERT) is based on a multi-layer bidirectional Transformer [21] and is trained on large plain texts for masked word prediction and next sentence prediction tasks.

PubTator [22] and BEST [23] are currently the two most comprehensive web platforms that can automatically mine compounds and (target) proteins from PubMed or PubMed Central (PMC). However, these tools fail to capture the compound-target relationships (interactions), and the resulting output may or may not contain experimental data. To solve these shortcomings, we set out to construct a pipeline using a BERT-based text classifier to identify studies containing DTIs and extract the associated data. We trained several BERT models (i.e., BERT, SciBERT [13], BioBERT [24], BioMed-RoBERTa [25] and BlueBERT [26]) on known studies containing DTIs and used majority voting of five BERT models to predict new studies containing DTIs. We identified approximately 2.1M new studies that possibly have DTI in terms of quantitative bioactivities. The identified studies are further linked with mined compound and protein entities provided by PubTator. Furthermore, the proposed BERT-based model could predict the assay format used in the experiment with an accuracy of ~90%. The resulting predicted and integrated data is freely available at https://dataset.drugtargetcommons.org/.

## MATERIALS AND METHODS

### Compound and protein annotations for studies on PubMed

We downloaded compounds and proteins identified from the full text of 24M documents (75% of PubMed) using PubTator’s API [22]. We define here document as a merged text containing titles and abstracts for the studies. We extracted data in batches of 1000 documents using parallel requests to save time. After preprocessing, we truncated those documents where text size exceeds 512 words because the sequence limit for BERT tokenization is 512. Approximately a quarter of the studies at PubTator missed the abstract information. We considered only those studies for which both abstract and title information is present in PubTator and therefore left with ~18.5M documents.

### Known studies for compound-target bioactivity data

Data used for the model’s training contains 66,521 positive examples (studies containing compound-target bioactivity data) and 66,521 negative examples (other biological studies), as shown in Supplementary File 1. Compound-target studies are extracted from DrugTargetCommons and ChEMBL (27^th^ release), whereas data for other biological documents is extracted from DisGeNET [27]. DisGeNET is a comprehensive resource for cataloguing gene-disease associations. Abstracts and titles (for studies) are extracted using PubMed’s API. Trained models are then used to predict documents (abstract + title) likely to contain DTIs. Positively predicted documents are associated with compound and protein entities as identified by PubTator.

### Assay formats for compound-target bioactivity data

Furthermore, we trained our models to predict the assay format most likely used in the documents. Assay format annotations are extracted from DrugTargetCommons for 28,102 documents with 14,109 focusing on cell-based assays and 13,993 having other assay formats (e.g., biochemical (93), cell-free (66), organism-based (12,845), tissue-based (424) and physiochemical (565)). We merged several assay formats under the ‘other’ category to avoid data imbalance problems while predicting assay format as shown in Supplementary File 2.

### Proposed method

BERT base is a masked language model (MLM) with 12 layers of architecture, pre-trained on > 2.5B words from English Wikipedia. We used BERT base and other BERT models (SciBERT, BioBERT, BioMed-RoBERTa and BlueBERT) to identify new documents on PubMed containing DTIs. The five BERT models are merged using majority voting to predict the label for the documents. Majority voting is a technique in machine learning used to combine the prediction power of several models. We adapted the majority voting technique to have more confidence in the prediction of true labels. SciBERT is an MLM pre-trained model trained on 1.14M full-texts from Semantic Scholar corpus with 82% from the biomedical domain [28]. SciBERT uses a different vocabulary (SCIVOCAB), whereas BERT, in general, is based on BASEVOCAB. In this study, we adapted uncased SciBERT. BioBERT is an MLM pre-trained language model based on the BERT representation for the biomedical domain. We used BioBERT-v1.1, pre-trained on PubMed for 200K steps and 270K steps on PMC. The model is pre-trained using the same hyper-parameter settings as for the original BERT model. BioMed-RoBERTa is MLM pre-trained language model based on the RoBERTa [25]. Finally, BlueBERT is pre-trained on ~4B words extracted from PubMed.

We used the BERT representations for the classification task by fine-tuning the BERT variants with minimal changes applied during the training phase. All BERT models used in this analysis comprised 12 layers of transformer encoder with hidden state dimensions equal to 768 and having >110M parameters as adopted in [21]. In our architecture, we used the last hidden layer as the output layer, which is used to predict whether if the document is drug-target like or not. To get the representation of the text sequence with length 512 tokens (words), we have used the BERT special [CLS] token, which is used for classification and contains the embedding for the text sequence. The special [SEP] token is used to separate the text sequences. We then used 2 fully connected layers and finally applied SoftMax. The BERT variants are fine-tuned using NVIDIA Tesla V100 SXM2 32 GB GPU, with a batch size of 32, maximum sequence length: 512, a learning rate of 2e-5 and maximum epoch size of 3. We used Adam with β_1_=0.9 and β_2_=0.999, slanted triangular learning rates as in [18], warm-up portion to 0.1 and ensured that GPU memory is fully utilized. The model architecture for all BERT models in this study is shown in Figure 1.

Next, we divided the overall workflow into three modules:

1. To identify whether a study in PubMed is likely to contain bioactivity data for drug and protein pair or not.
2. Extract drug and protein information by taking advantage of already extracted entities by PubTator.
3. Predict assay format for positively identified studies.

**Figure 1:**
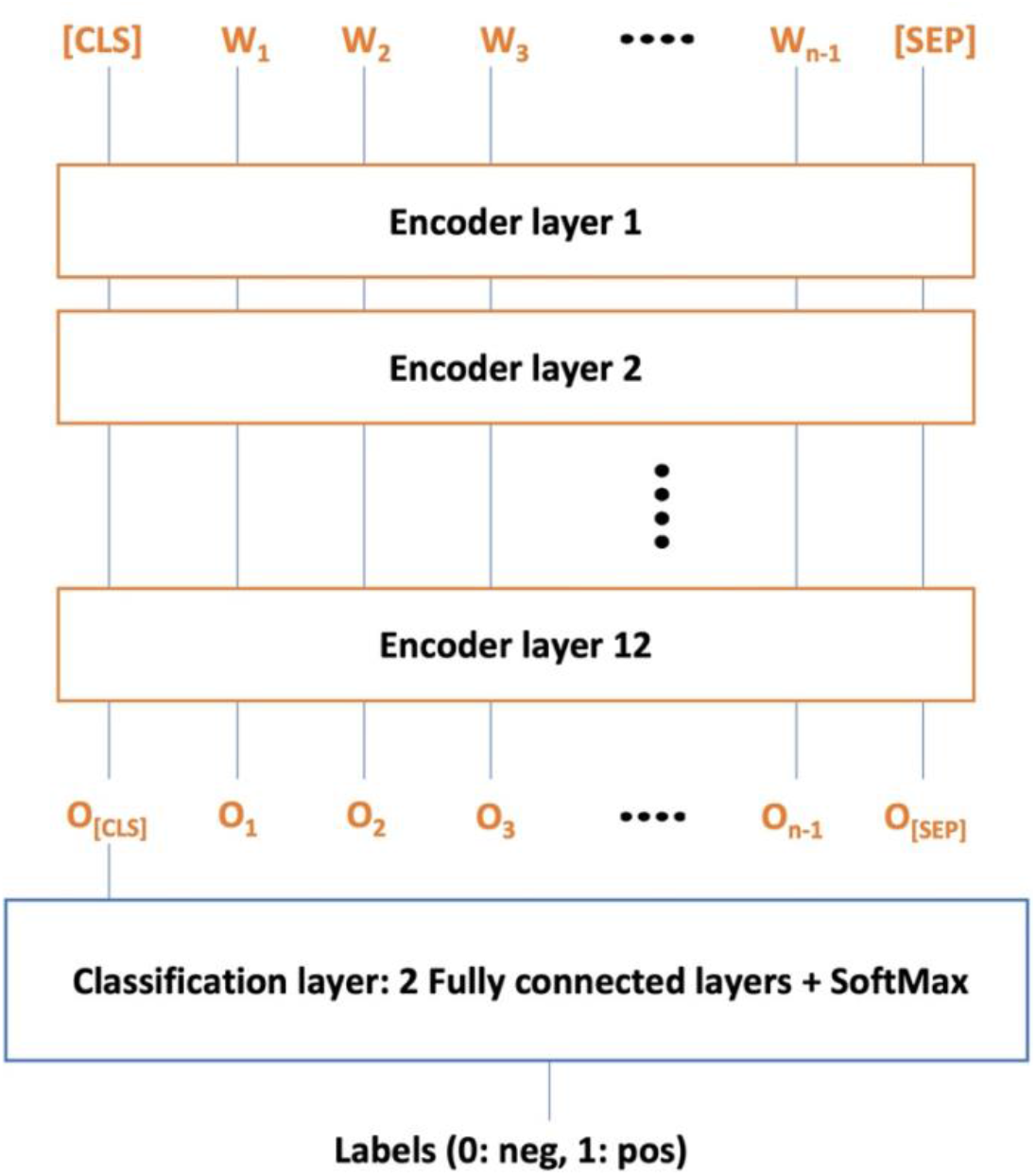
Architecture for all BERT variants, where Wi represents input word token and Oi represents contextual embeddings at the output layer. The O[CLS] is first token of output sequence and O[CLS] contains class label.

For module 1, we used the fine-tuned BERT models to predict whether PubMed study contains a compound-target relationship or not. BERT models are trained on 66,521 positive and 66,521 negative documents (abstracts + title of the study) as explained in the previous section. We pre-processed all the BERT models by assigning tokens for each word in the documents and converted all words into lower case. We then padded for cases where document length <512. Finally, each document (study) is mapped into 768 numeric features with minor differences in the five models. After training, BERT models are merged in majority voting to identify new studies possibly containing bioactivity DTIs. For module 2, we then matched and linked positively predicted documents with identified drug and gene entities using the PubTator dataset. Finally, for module 3, using the same architecture, we tried to predict assay formats for the positively predicted documents (cell-based vs other assays). We emphasized on assay format field because assay formats are critical in defining scores for DTIs [29]. We organized the final output in terms of PubMed id for study, predicted assay format, drugs, and proteins, and which is freely available at https://dataset.drugtargetcommons.org/. The workflow for the proposed research is shown in Figure 2.

**Figure 2:**
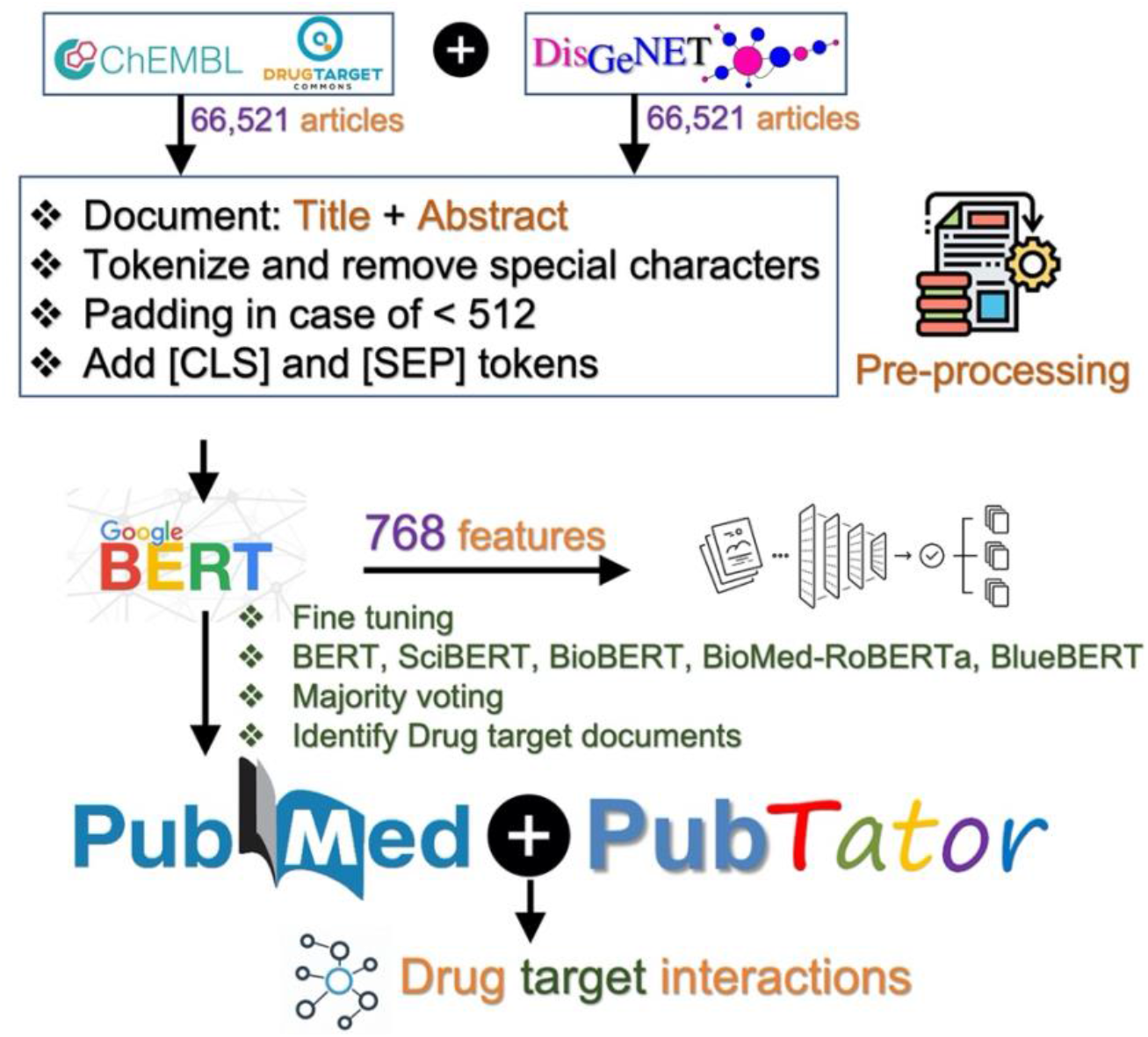
Workflow for identifying new studies containing drug-target bioactivity data. The drug and protein entities for studies are integrated from the PubTator dataset. The final output contains predictions for ~18.5M studies that possibly have DTIs

## RESULTS AND DISCUSSIONS

### Ten-fold cross-validation results using BERT models

To feed the PubMed documents into the BERT pipeline, we first converted linguistic units (text snippets, words and phrases that carry meaning) in 66,521 positive and 66,521 negative documents into tokens. We truncated those documents for which document length >512 (maximum limit by BERT). Each BERT model generated a numeric feature set of length 768 for each document. BERT text classifier is trained using 10-fold cross-validation, and the performances are shown in Table 1.

**Table 1:**
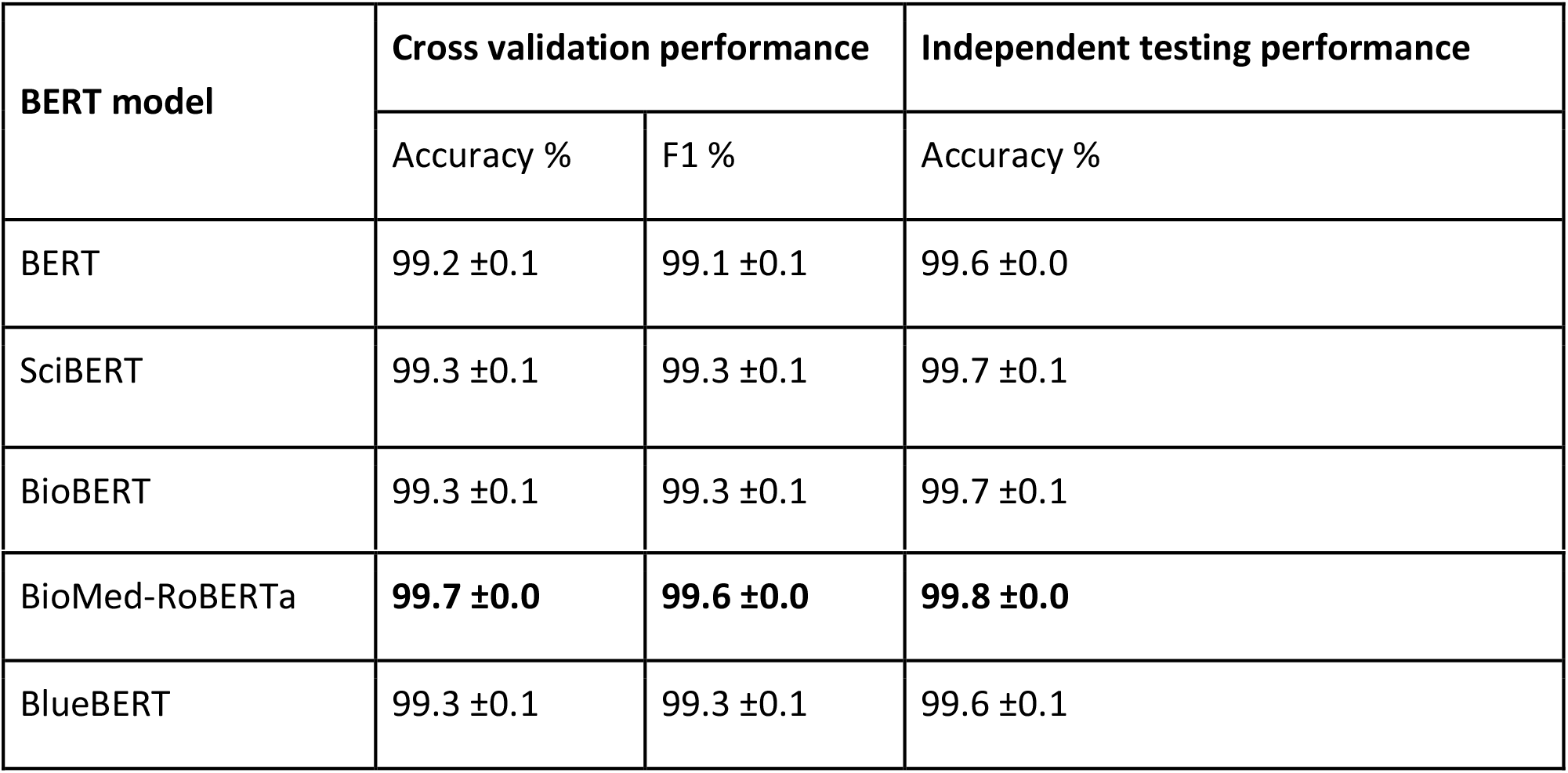
Ten-fold cross-validation results for different BERT models. The last column shows accuracy% for independent testing of negative class.

Our analyses showed that all BERT models reached accuracies higher than 99%. In comparison, the traditional machine learning algorithms including Support Vector Machine and Naïve Bayes yielded accuracies of 98.87% and 98.29%, respectively. The accuracy of independent BERT testing on 7,785 studies of only the negative class (other biological documents) also showed > 99% accuracy. Our findings demonstrate that BERT models can successfully detect drug-target like documents and distinguish true negatives with great precision.

To examine the word distributions in two types of documents, we also analyzed the top frequently occurring words in positive and negative documents. As shown in Figure 3, the most frequently occurring words in drug target documents are ‘compounds’, ‘activity’ and ‘potent’, whereas the most frequent words for other biological documents are ‘expression’, ‘patients’, and ‘gene’. Word distribution analysis can demonstrate developing a simple model based on word frequencies to identify drug-target or other biological documents.

**Figure 3:**
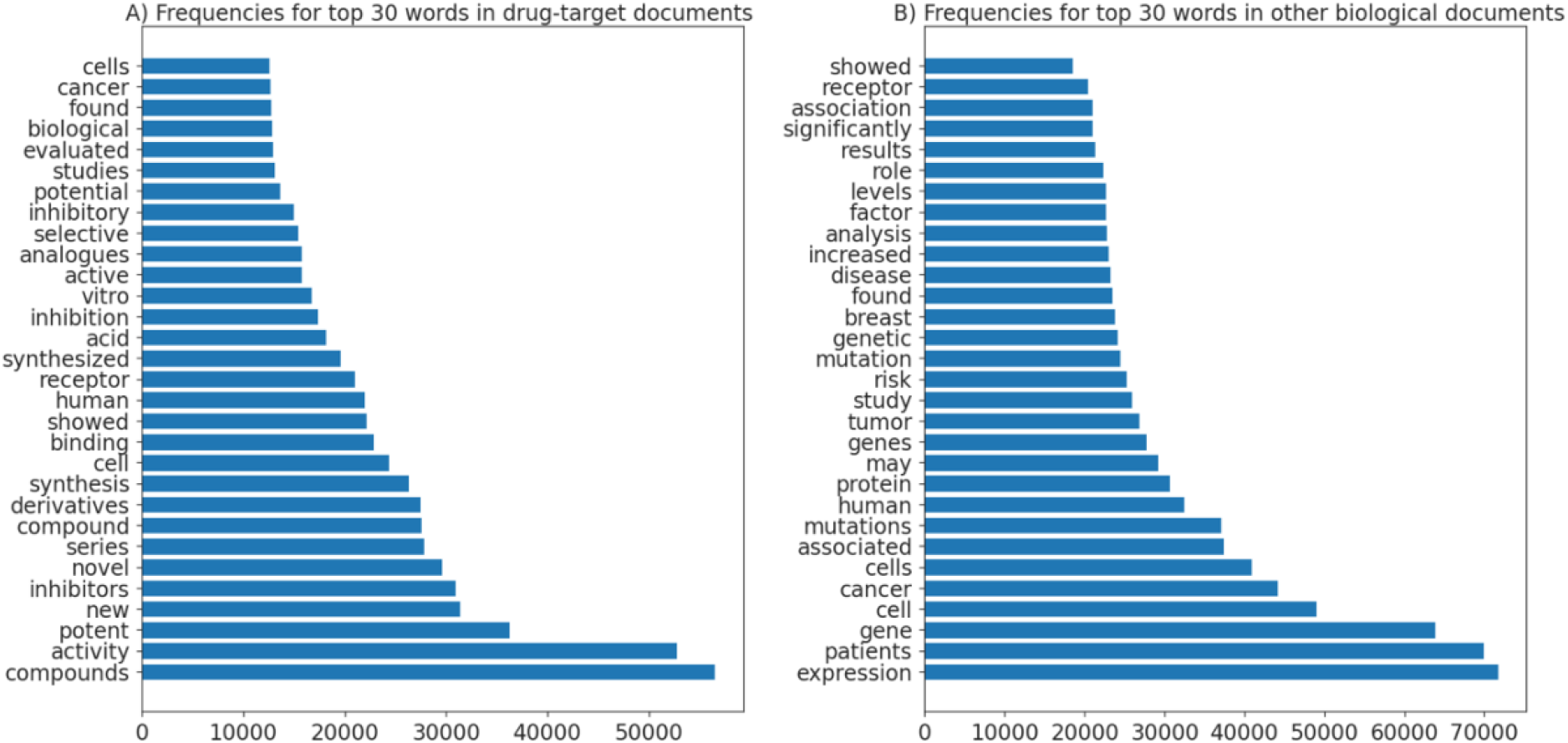
Top 30 words frequencies, A) Drug-target like documents, B) Other biological documents.

### Identify new drug-target studies and associate drug and protein pairs using PubTator dataset

After successfully training the BERT models, we tried to identify new studies on PubMed that possibly contain bioactivity data for drug-target pairs. For this purpose, we used 18.5M documents downloaded using PubTator’s API. Each BERT model has its strength, and we used the predictions from the five BERT models in majority voting to determine whether a study classification is positive or negative. Table 2 shows the number of positively and negatively predicted documents out of 18.5M documents. The fourth column (Studies containing drugs or proteins on PubTator) in Table 2 shows how many among positively predicted studies (using BERT models) have either drug or protein entity extracted by PubTator. Finally, the last column indicates the number of studies for which PubTator identified both drugs and proteins. These two columns, validate those studies that are identified as drug-target studies using BERT model.

**Table 2:** Prediction of drug-target like documents from 18.5M studies on PubMed. The fourth column shows the number of documents that contain either drug or gene entities as identified by PubTator. In contrast, the fifth column indicates the number of documents that contain both drugs and genes entities.

| BERT model | Predicted as drug-target studies | Predicted as other studies | Studies containing drugs or proteins on PubTator | Studies containing both drugs and proteins on PubTator |
| --- | --- | --- | --- | --- |
| BERT | 2,394,207 | 16,164,382 | 2,239,986 | 513,972 |
| SciBERT | **2,564,440** | 15,994,149 | **2,386,665** | **562,051** |
| BioBERT | 1,914,147 | 16,644,442 | 1,821,758 | 459,349 |
| BioMed-RoBERTa | 1,371,113 | 17,187,476 | 1,311,936 | 315,756 |
| BlueBERT | 2,443,832 | 16,114,757 | 2,304,304 | 530,033 |
| Majority voting | 2,129,731 | 16,428,858 | 2,019,050 | 467,638 |

Superior performance on unseen studies shows how well the predictions by BERT models can be generalized. Using the majority voting of BERT models, 94.8% (2,019,050) of the studies identified as drug-target like (positive) containing either drugs or proteins entity identified by PubTator. Out of the ~2.1M positively predicted documents, 21.9% (467,638) contain both drug and protein entities at PubTator. The result is likely an underestimation, as drug or protein entities (or both) may have been deposited as supplementary data, which is not captured by PubTator’s back-end algorithm. This means that even though the study is positively predicted our workflow might not capture drugs or proteins in some cases, leaving the task for manual curators to check the supplementary material. Indeed, many high throughput studies do not mention drug or protein names in the article’s main text but instead provide these in the supplement (e.g. Davis et al., 2011) [31]. Of the BERT models, SciBERT identified more drug-target like documents compared to other models, with at least 562,051 studies containing both drugs and proteins in the PubTator dataset. This could be because SciBERT is additionally trained for biomedical applications, whereas other models are designed for general text classification applications.

We also analyzed the top journals, and yearly distribution for ~2.1M predicted studies. This will give an idea for manual curators as to what journals (and year of publications) are suitable for extracting DTIs. Using the Bio Entrez package in python, we obtained journal names and year of publication for ~2.1M studies. We extracted journal names and year information only for ~0.2M out of ~2.1M studies (though PubMed IDs are present in Entrez). Journal names and years of publications for most of the studies are missing from the Entrez database. However, we still can have an idea of the general trend based on captured information. Figure 4A shows the top 15 journals containing drug-target like studies, whereas Figure 4B shows year wise frequencies for the studies. As shown in Figure 5A, Journal of Medicinal Chemistry, Biological Chemistry, and Bioorganic & Medicinal Chemistry Letters are present among the top 15 journals. These three journals are among the leading journals for bioactivity data extraction in ChEMBL [32]. Furthermore, most drug-target studies are from recent years, with the year of 2020 containing the most significant number of studies.

**Figure 4:**
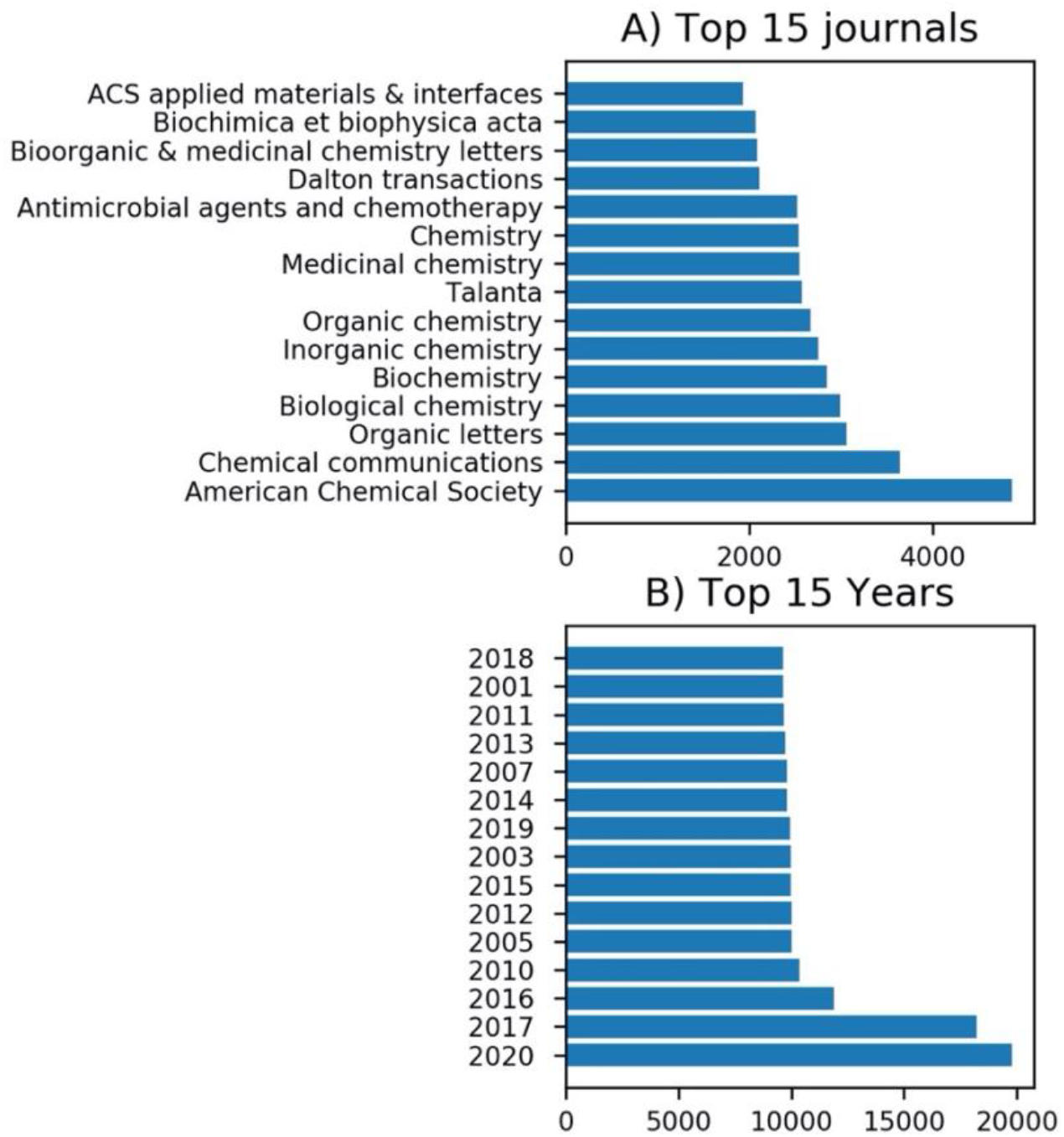
Statistics for ~0.2M studies which are predicted as drug-target like articles using majority voting, A) Top 15 journals and B) Top 15 years for drug-target like studies.

**Figure 5:**
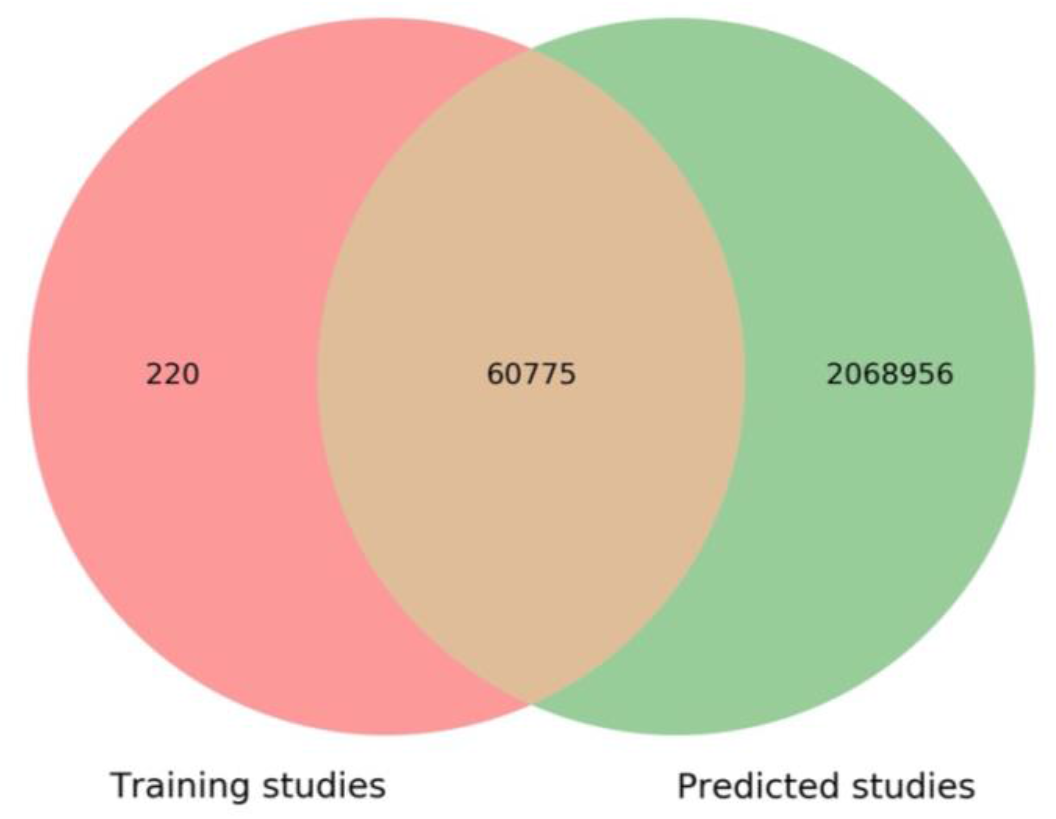
Overlap between training and predicted studies.

There are 66,521 DTI studies that are used to train BERT models. Among these, 60995 are overlapping with 18.6M studies in PubTator dataset. We also tried to analyze the overlap of 60,995 DTI studies with ~2.1M studies that are predicted as DTI studies using BERT models. Figure 5 shows that 99.6% (60,775) DTI studies are present among the list of predicted DTI studies. This means our analysis identified 2,068,956 additional studies containing DTIs. These newly identified studies, along with PubMed IDs, drugs and proteins involved in the study, are freely available at https://dataset.drugtargetcommons.org/. The output of our analysis can be used as a starting point to further extract the quantitative drug-target bioactivity values from the identified studies. We hope that our output will significantly ease the job of manual curators as we are providing the actual PubMed ID, drugs, and protein entities, as well as assay formats for ~2.1M identified studies.

### Predict assay format for drug-target studies

After successfully identifying DTIs, the next task is to predict the assay format most likely adapted in the identified studies. For that purpose, we separately trained each BERT model on assay format dataset with 14,109 documents based on cell-based and 13,993 having other assay formats. We used the same fine-tuning settings as for drug-target study identification task. Figure 6 shows 10-fold cross-validation performances for BERT models in terms of accuracy% and F1%. All BERT models have accuracy >88% with BioBERT slightly outperforming other models. This shows that BERT models can successfully identify drug-target studies as well as assay formats.

**Figure 6:**
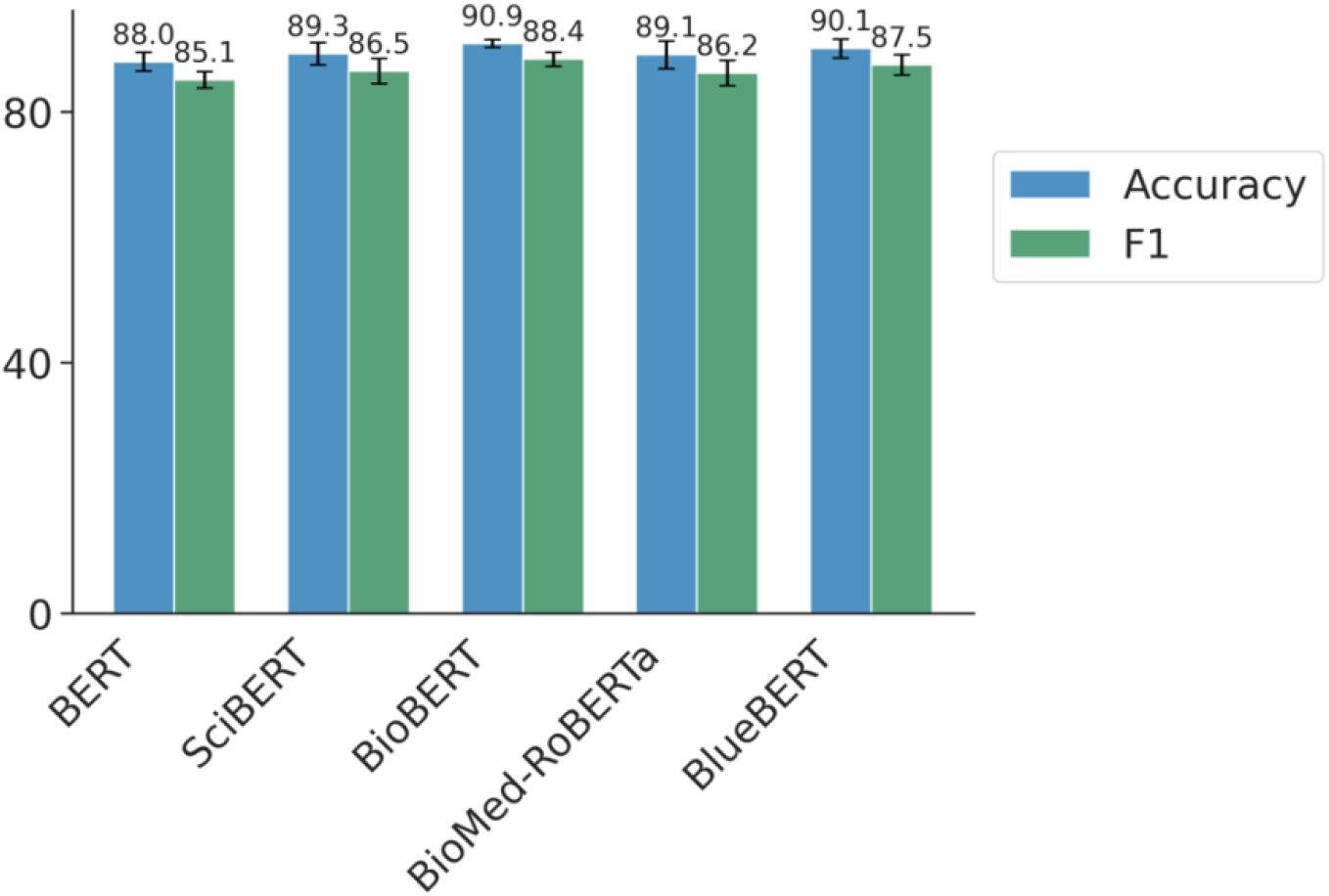
10-fold cross-validation performance for the task of assay format prediction.

After successful training of BERT models to predict assay formats (on known studies), we applied trained models for predicting assay formats in those studies, which are positively predicted using majority voting. There are ~2.1M studies that are predicted as drug-target like document using majority voting as mentioned in Table 2 (last row). Therefore, we tried to predict assay formats for those ~2.1M studies. As shown in Table 4, >26.5% of the drug-target like studies (564,425 out of 2,129,731) are likely based on cell-based assays. It is impossible to computationally validate these predictions now. However, in the next release of DrugTargetCommons, we will validate the predicted assay formats using manual curation of ~2.1M identified studies.

**Table 4:** BERT models to predict assay formats for ~2.1M newly predicted drug-target like studies.

| BERT models | Cell based assay | Another assay |
| --- | --- | --- |
| BERT | 864,319 | 1,265,412 |
| SciBERT | 541,322 | 1,588,409 |
| BioBERT | 474,283 | 1,655,448 |
| BioMed-RoBERTa | 1,013,497 | 1, 6,234 |
| BlueBERT | 476,716 | 1,653,015 |
| Majority voting | 564,425 | 1,565,306 |

## CONCLUSIONS

Most DTI resources are compound centric and lack DTI profiles at complete proteomic level. In compound centric approaches, thousands of the compounds are tested across a particular target protein. Only 11% of the human proteome are targeted by small molecules or drugs, whereas one in three proteins is still being investigated [33]. The combined non-overlapping experimental studies (on PubMed) are less than 0.1M. Curating quantitative bioactivity values reported in a study cannot be fully automated. However, semi-automated NLP based methods can assist in identifying related studies and easing the workload for the manual curators. BERT is recently proposed as a state-of-the-art model for several NLP tasks, including text classification. Therefore, in this research, we investigated several models of BERT to identify new studies possibly containing DTIs.

Furthermore, we tried to predict assay formats most likely used in the studies. Assays formats, along with actual bioactivities, are critical in defining scores for DTIs. Using majority voting based on BERT models, we identified 2,129,731 studies from which 467,638 are confirmed to have both drug and protein entities in the PubTator dataset. Most of these ~2.1M studies are not reported in any of the manually curated bioactivity databases as the combined non-overlapping studies curated by commonly used DTI databases are around 0.1M. These identified DTIs (along with annotations) are freely available at https://dataset.drugtargetcommons.org/. We hope that the identified studies and drug and protein entities will ease the job of manual curators and improve protein target coverage across investigational and approved compounds. Lastly, increased target coverage for investigational and approved will enhance the understanding of drug mechanism of action and open new drug repurposing opportunities. The manual curation team of DrugTargetCommons will take advantage of newly identified studies and curate more bioactivity data in their next release.

## DATA AVAILABILITY STATEMENT

Newly identified studies extracted drug/gene entities and predicted assay formats are freely available at https://dataset.drugtargetcommons.org/.

